# Neurobiological underpinnings of rapid white matter plasticity during intensive reading instruction

**DOI:** 10.1101/2020.05.28.122499

**Authors:** Elizabeth Huber, Aviv Mezer, Jason D. Yeatman

## Abstract

Diffusion MRI is a powerful tool for imaging brain structure, but it is challenging to discern the biological underpinnings of plasticity inferred from these and other non-invasive MR measurements. Biophysical modeling of the diffusion signal aims to render a more biologically rich image of tissue microstructure, but the application of these models comes with important caveats. A separate approach for gaining biological specificity has been to seek converging evidence from multi-modal datasets. Here we use metrics derived from diffusion kurtosis imaging (DKI) and the white matter tract integrity (WMTI) model along with quantitative MRI measurements of T1 relaxation to characterize changes throughout the white matter during an 8-week, intensive reading intervention (160 total hours of instruction). Behavioral measures, multi-shell diffusion MRI data, and quantitative T1 data were collected at regular intervals during the intervention in a group of 33 children with reading difficulties (7-12 years old), and over the same period in an age-matched non-intervention control group. Throughout the white matter, mean ‘extra-axonal’ diffusivity was inversely related to intervention time. In contrast, model estimated axonal water fraction (AWF), overall diffusion kurtosis, and T1 relaxation time showed no significant change over the intervention period. Both diffusion and quantitative T1 based metrics were correlated with pre-intervention reading performance, albeit with distinct anatomical distributions. These results are consistent with the view that rapid changes in diffusion properties reflect phenomena other than widespread changes in myelin density. We discuss this result in light of recent work highlighting non-axonal factors in experience-dependent plasticity and learning.

**Highlights:**

- **Diffusion MRI measurements in white matter show changes linked to an educational intervention.**
- **Tissue modeling results point to changes within the extra-axonal space.**
- **Complementary MRI measurements fail to suggest a widespread change in white matter in myelination over the intervention period.**
- **Both diffusion and quantitative T1 measures correlate with pre-intervention reading skill.**

## Introduction

Experience can alter the structure of white matter throughout life (Bechler et al., 2018; Fields, 2015; Sampaio-Baptista and Johansen-Berg, 2017). In human subjects, evidence for white matter plasticity has been reported after remarkably brief behavioral interventions, ranging in length from hours to months (Ekerdt et al., 2020; Hofstetter et al., 2017; Hofstetter et al., 2013; Huber et al., 2018; Jolles et al., 2016; Mamiya et al., 2016; Metzler-Baddeley et al., 2017; Sampaio-Baptista et al., 2013). Experience dependent plasticity is therefore thought to occur not only during specific developmental windows, but well into adulthood (see Sampaio-Baptista and Johansen-Berg, 2017 for a recent review). However, it is challenging to determine the neurobiological underpinnings of these effects: For one, the majority of past work has relied on diffusion MRI (dMRI) and scalars derived from the diffusion tensor model (DTI, Basser et al., 1994), which entangle microstructural features with the spatial organization of axon bundles at the voxel level, diminishing microstructural specificity (De Santis et al., 2014). Broadly speaking, parametric variation of one kind (e.g., radial diffusivity) does not map onto specific variation in tissue structure (e.g., myelination). The same is true for diffusion kurtosis imaging (DKI, Jensen et al., 2005), which accounts for higher order variation in the diffusion signal and extends the tensor model to larger b-values (>1000 s/mm^2^). Both DTI and DKI provide descriptions of the diffusion signal that are agnostic to tissue composition. Modeling approaches that offer a biologically richer interpretation of the diffusion signal thus hold an appeal.

Over the last two decades, a host of biophysical models have been proposed with the aim of better characterizing the diffusion signal with respect to underlying tissue structure (Alexander et al., 2019; Jelescu and Budde, 2017). In general, these models envision two or more non-exchanging, tissue-specific water compartments (e.g., intra-versus extra-axonal) whose distinct signal decay profiles can be leveraged to yield relative weightings (signal fractions) for each compartment. For example, DKI based tissue models have been used to derive voxel-level neurite/axon fractions and compartment specific intra- and extra-axonal tensors (Fieremans et al., 2011; Jensen and Helpern, 2010; Jespersen et al., 2010). A recent implementation (Fieremans et al., 2011), known as the ‘white matter tract integrity’ (WMTI) model (Benitez et al., 2014), has been used to characterize axonal pathology and de-myelination (Falangola et al., 2014; Guglielmetti et al., 2016; Jelescu et al., 2015; Jelescu et al., 2016b; Kelm et al., 2016). WMTI parameters have also been linked to individual differences in cognitive performance in healthy adults (Chung et al., 2018a) and children (Huber et al., 2019), as well as variation in the white matter as a function of age during early development (0-3 years, Jelescu et al., 2015), throughout childhood and adolescence (4-19 years, Genc et al., 2017), and across the lifespan (7-63 years, Chang et al., 2015). Indeed, with appropriate pre-processing, these parameters are highly reliable, even in children, for whom head motion is a particular concern and scan durations are necessarily limited (Huber et al., 2019).

As is true for any modeling framework, the interpretability of results depends strongly on the validity of underlying assumptions. In the WMTI model, intra- and extra-axonal compartment specific tensors are derived in light of the axonal water fraction (AWF) estimated voxel-wise from the overall kurtosis tensor (Fieremans et al., 2011). For complex fiber configurations, AWF estimates are approximate and provide theoretical upper/lower limits on compartment fractions, absent a priori knowledge of the intra-axonal diffusivity (Fieremans et al., 2011; Jespersen et al., 2010). Although originally developed in the context of aligned fibers, it is possible to derive AWF, along with intra- and extra-axonal compartment diffusivities, for arbitrary configurations of fibers: One can proceed by factoring out orientation dispersion via directional averaging (Henriques Rafael Neto, 2021; Kaden et al., 2016), by using a pre-defined ODF (e.g., Jespersen et al., 2018; Zhang et al., 2012), or by incorporating a data-derived estimate of the orientation distribution function (ODF) (see Novikov et al., 2018; see Tournier, 2019 for recent review of approaches). Recently, McKinnon and colleagues (2018) used ODF estimates from fiber ball imaging (FBI) (Jensen et al., 2016) to inform estimates of AWF and compartmental diffusivities throughout the white matter, providing a direct extension of WMTI to white matter regions with more complex architecture. Importantly, McKinnon and colleagues (2018) also demonstrated discrepancies in intra-axonal diffusivity estimates across models with and without the FBI-derived ODF, particularly in regions with lower FA (<0.8). Other recent work has also highlighted the confounding of compartment diffusivities, signal fractions, and dispersion values, in the context of a model in which all parameters are freely fit (Jelescu et al., 2016a). An ensuing debate as to the appropriate designation of ‘faster’ vs. ‘slower’ diffusing compartments in the WMTI model (see Jelescu et al., 2020 for recent review) serves to underscore present challenges for relating diffusion measurements to underlying tissue, and the need for caution in assigning a biological interpretation to diffusion parameters.

Although tissue modeling techniques may yield ambiguous or even inaccurate results, model parameters may still help to inform new linking hypotheses, in conjunction with complementary imaging measures, such as quantitative estimates of T1 relaxation and magnetization transfer (MT). Multi-modal studies that combine diffusion MRI with these measures can provide critical insight (Cercignani and Bouyagoub, 2018; Filo et al., 2019; Takemura et al., 2019; Travis et al., 2019). In a recent study with pre-school aged children, T1 relaxation, but not FA, in the anterior arcuate fasciculus predicted family history of dyslexia and subsequent childhood reading performance (Kraft et al., 2016), suggesting that behaviorally important differences in myelination might be obscured by the lack of microstructural specificity inherent in the diffusion metrics. At the same time, a number of studies have reported developmental changes in diffusion properties over adolescence (Lebel and Beaulieu, 2011; Tamnes et al., 2018), while recent work has failed to detect a corresponding change MT based metrics (Moura et al., 2016). In a large cross-sectional study, complementary DTI and quantitative T1 relaxation measurements showed distinct trajectories across age (Yeatman et al., 2014). Recent multi-modal analyses have sought to disentangle effects attributed to factors like axonal size and packing versus myelination (Geeraert et al., 2019), potentially offering a more nuanced view of white matter development. Where discrepancies across imaging modalities stem from differential sensitivity to specific tissue features, rather than differences in signal to noise (either intrinsically, or by virtue of methodological factors yielding SNR differences across data types), they may be highly informative about biological processes unfolding with distinct time courses over development, or with distinct roles in predicting behavior.

We previously reported that 8 weeks of intensive, native-language (English) reading instruction prompts widespread changes in white matter diffusion properties, which track the learning process (Huber et al., 2018). Given that these effects occurred rapidly, and were distributed throughout the white matter, we speculated that the changes in diffusivity might reflect an initial stage of the learning process, rather than functionally specific remodeling of myelin. Here, we first re-examined the previously described intervention effect in a larger sample of participants (n=32). We then tested for longitudinal changes in parameters derived from the WMTI model, as well as in quantitative R1 relaxation rates (1/T1 relaxation time). At baseline (pre-intervention), diffusion parameters and R1 estimates both correlated with reading skill; however, the anatomical distribution of these effects differed across modalities. Although we observed compartment specific changes in diffusivity, we failed to detect a change in AWF or R1.

## Materials and Methods

### Participants

A total of 128 behavioral and MRI sessions were conducted with a group of 34 children ranging in age from 7 to 12 years, who participated in an intensive summer reading intervention program. Of these participants, 24 were included in a previous manuscript (Huber et al., 2018). Members of the intervention group were recruited based on parent report of reading difficulties and/or a clinical diagnosis of dyslexia. Multi-shell diffusion MRI and behavioral data were collected before the intervention (baseline), after 3.63 weeks of intervention (+/− 1.49), after 6.77 weeks of intervention (+/− 1.32), and at the end of the 8-week intervention period. An additional 87 behavioral and MRI sessions were conducted with 45 participants, who were recruited as a control group to assess the stability of our measurements over the repeated sessions. A total of 29 sessions (out of 215 total) were removed from further analysis due to excessive head motion or other artifacts in the MRI data, as described below. The final sample included 111 sessions from 32 intervention participants (12 female, mean age 9.43 years), and 75 sessions from 41 control participants (16 female, mean age 9.86 years). The final control group was matched in age (*t*(71) = −1.17, *p* = 0.25) but not reading level. Pre-intervention behavioral scores for both groups are shown in **Figure 1**.

**Figure 1.**
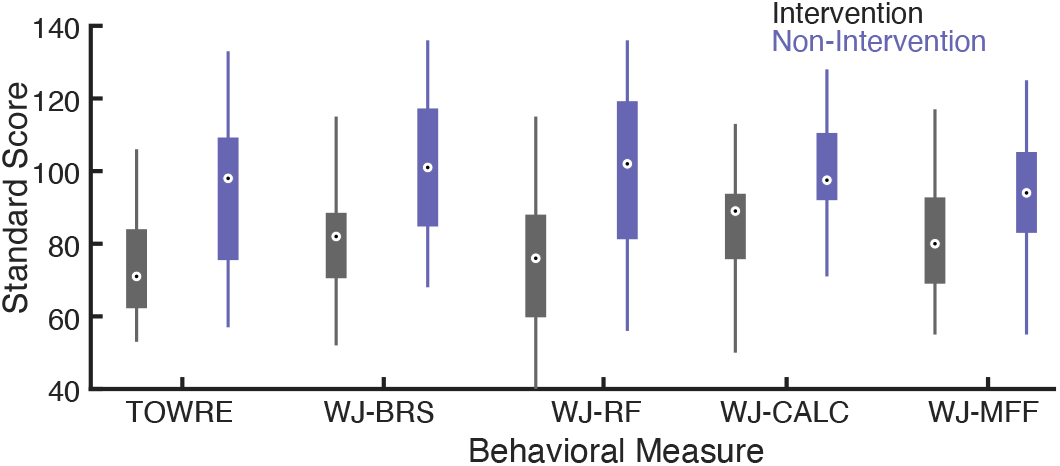
Boxplots of pre-intervention period reading and math scores. Median scores (open circles) are plotted for the intervention (gray) and non-intervention control (blue) groups. Box edges mark the 25th and 75th percentiles, while whiskers span the full range of data points for each measure: Test of Word Reading Efficiency index (TOWRE) and the Woodcock Johnson Tests of Achievement subtests for Basic Reading Skill (WJ-BRS), Reading Fluency (WJ-RF), Calculation (WJ-CALC), and Math Facts Fluency (WJ-MFF).

Control participants completed the same experimental sessions but did not receive the reading intervention. Some families of control group participants were reluctant to commit to all four sessions, given that their children were not receiving an educational intervention. These families were given an opportunity to participate in 2 sessions. The interval for two sessions was chosen to produce balanced numbers of measurements at equivalent time points to the intervention group. In the intervention group, 5 participants were unable to complete all imaging sessions, and therefore participated in 3 sessions, total. One intervention participant who completed only a single MRI session, for scheduling reasons, was excluded from the longitudinal analysis as well as cross-sectional analyses, based on data quality. The distribution of testing sessions for the final sample of intervention and control participants, after excluding subjects for quality control (e.g., due to excessive motion during MRI), is summarized in **Figure 2**.

**Figure 2.**
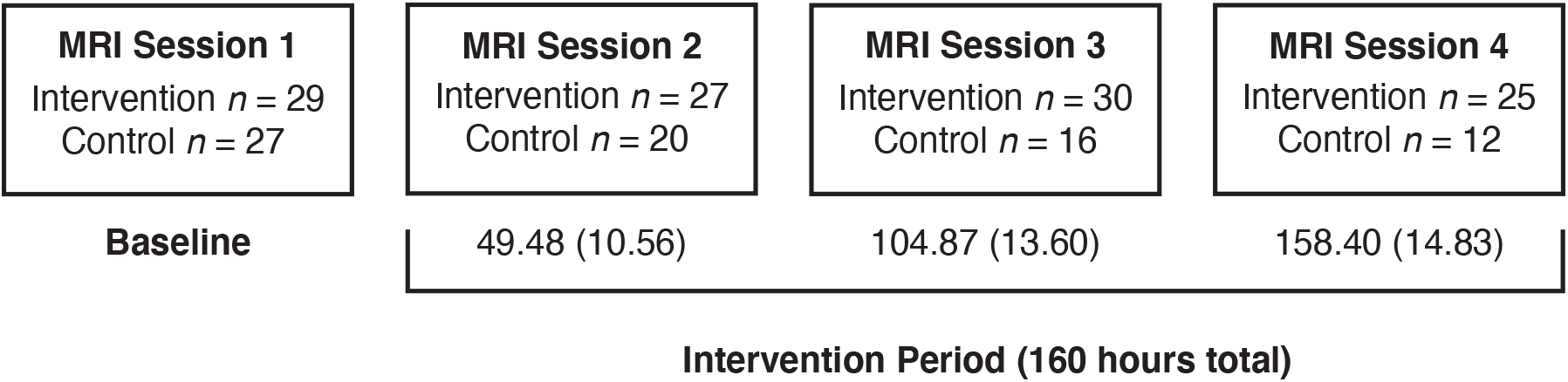
Testing schedule for intervention and control groups. Experimental sessions were evenly spaced for each participant, with a baseline prior to the start of intervention, and three timepoints during the intervention period, after approximately 3, 6, and 8 weeks of instruction. The number of participants in the intervention and control groups that survived data quality control procedure is given for each time point, along with the mean number of instruction hours completed by the intervention group, with standard deviation in parentheses. Intervention participants completed a total of 160 hours of reading instruction.

All participants were native English speakers with normal or corrected-to-normal vision and no history of neurological damage or psychiatric disorder. Participants first completed a mock scan to familiarize them with the scanning environment and to ensure their comfort and ability to hold still during the MRI sessions. We obtained written consent from parents, and verbal assent from all child participants. All procedures, including recruitment, consent, and testing, followed the guidelines of the University of Washington Human Subjects Division and were reviewed and approved by the UW Institutional Review Board.

### Reading intervention

Intervention participants were enrolled in 8 weeks of the Seeing Stars: Symbol Imagery for Fluency, Orthography, Sight Words, and Spelling (Bell, 2007) program at three different Lindamood-Bell Learning Centers in the Seattle area. The intervention program consists of directed, one-on-one training in phonological and orthographic processing skills, lasting four hours each day, five days a week. The curriculum uses an incremental approach, building from letters and syllables to words and connected texts, emphasizing phonological decoding skills as a foundation for spelling and comprehension. A hallmark of this intervention program is the intensity of the training protocol (4 hours a day, 5 days a week) and the personalized approach that comes with one-on-one instruction.

To test for longitudinal change in reading and non-reading (Calculation and Math Facts Fluency subtests of the Woodcock Johnson Tests of Achievement) measures, we fit a linear mixed effects model with a fixed effect of intervention time, in days (for control participants, this corresponds to days since the baseline session), and a random effect of participant (random intercept). All statistical analyses were carried out using custom software written in MATLAB (version R2018a) and the MathWorks Statistics Toolbox. The intervention group showed significant gains for all reading measures (p < 0.0001). We found no significant growth in calculation (*t*(96) = −1.85, *p* = 0.068) or math facts fluency (*t*(101) = −1.80, *p* = 0.074), both of which decreased marginally over the same time period. Note that the degrees of freedom above reflect occasional missing behavioral data due to participant fatigue in a small number of sessions. In the control group, performance on timed reading and non-reading measures (TOWRE, WJ-RF, and WJ-MFF) improved with repeated testing (TOWRE: *t*(73) = 3.89, *p* = 0.00022, WJ-RF: *t*(71) = 3.32, *p* = 0.0014, WJ-MFF: *t*(73) = 2.22, *p* = 0.030), although reading accuracy and calculation showed no significant change (WJ-BRS: *t*(73) = 1.66, *p* = 0.10, WJ-CALC: *t*(70) = 1.52, *p* = 0.13). As above, degrees of freedom above reflect occasional missing behavioral data (for WJ-RF and WJ-CALC) due to participant fatigue. In a subset of reading matched controls (defined as having a basic reading WJ-BRS or timed reading TOWRE score 2 or more standard deviations below the population mean at Session 1), we saw an increase in reading fluency with repeated testing (WJ-RF: *t*(15) = 2.77, *p* = 0.014), and no significant change in any other measure.

### Magnetic resonance imaging (MRI) acquisition protocol

All imaging data were acquired using a 3T Phillips Achieva scanner (Philips, Eindhoven, Netherlands) at the University of Washington Diagnostic Imaging Sciences Center (DISC) using a 32-channel head coil. An inflatable cap minimized head motion, and participants were continuously monitored through a closed-circuit camera system.

Diffusion-weighted magnetic resonance imaging (dMRI) data were acquired at 2.0mm isotropic spatial resolution with full brain coverage. Each session consisted of 3 DWI scans, one with 32 non-collinear directions (b-value=800s/mm^2^), and a second with 64 non-collinear directions (b-value=2,000s/mm^2^). Each of the DWI scans included 4 volumes without diffusion weighting (b-value=0). We also collected one scan with 6 non-diffusion-weighted volumes and a reversed phase encoding direction (posterior-anterior) to correct for EPI distortions due to inhomogeneities in the magnetic field using FSL’s topup (version 5.0.9) tool (Andersson et al., 2003). Additional pre-processing was carried out using the FSL Eddy tool (Andersson and Sotiropoulos, 2016) for motion and eddy current correction. Diffusion weighted volumes were aligned first to the average of the non-diffusion weighted volumes, and then to an ACPC aligned T1 weighted anatomical image (MPRAGE acquired at 1.0mm isotropic) using a rigid body transformation (SPM version 12 Ashburner and Friston, 1997). Diffusion gradients were rotated to account for rotation applied during motion correction and alignment. Data were manually checked for imaging artifacts and excessive dropped volumes. Given that participant motion can be especially problematic for the interpretation of group differences in DWI data (Yendiki et al., 2014), data sets with mean slice-by-slice displacement > 3mm were automatically excluded from further analysis. In total, 14 data sets were excluded due to participant motion. In Session 1, two intervention participants and one control participant were excluded. In Session 2, four intervention participants and two control participants were excluded. In Session 3, one intervention and one control participant were excluded. In Session 4, three intervention participants were excluded.

For quantitative T1 mapping, we followed protocol developed by (Mezer et al., 2013). Briefly, we acquired 4 spoiled gradient echo recalled images using two different flip angles (2 scans with 4° and 2 scans with 20°, all with TR = 14ms, TE = 2.3ms, and resolution of 1mm^3^), which were aligned to the same T1 weighted anatomical image (MPRAGE acquired at 1.0mm isotropic) as the diffusion data (see above) using a rigid body transformation (SPM version 12 Ashburner and Friston, 1997). To correct the transmit coil inhomogeneity, we collected 4 spin echo inversion recovery scans with EPI read-out (SEIR-EPI), with TR 6500ms, TE 6.46ms, inversion times of 50, 400, 1200, and 2400ms, and 2mm^2^ in-plane resolution with a slice thickness of 4mm. The ANTs software package (Tustison et al., 2009) was used to register the spoiled gradient echo recalled images to the spin echo inversion recovery images. We then compared T1 fits estimated using the spoiled gradient echo images to fits estimated using the unbiased (Barral et al., 2010; Mezer et al., 2016; Mezer et al., 2013) SEIR images to characterize the inhomogeneity field and apply an appropriate correction to the biased, high resolution spoiled gradient echo recalled images using freely available, custom MATLAB code (https://github.com/mezera/mrQ). R1 maps were then created by taking 1/T1 (seconds) using the bias corrected T1 estimate for each voxel.

### Modeling white matter tissue properties

Axonal water fraction and compartment specific diffusivities were calculated according to the white matter tract integrity (WMTI) model as implemented in DIPY version 0.16.0 (Garyfallidis et al., 2014), after fitting the diffusion kurtosis model (Jensen et al., 2005) to the diffusion data, and following the pre-processing steps described above.

Modeling was carried out first with the default assumption of greater extra-versus intra-axonal diffusivity (Fieremans et al., 2011, see also). Specifically, extra-axonal diffusivity was defined as

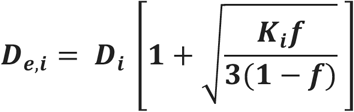

while intra-axonal diffusivity was defined as

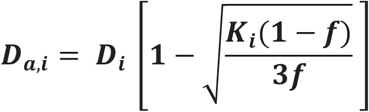

where *D_i_* and *K_i_* are the diffusion and kurtosis coefficients for *i* = 1,2,3, corresponding to the axes of the *overall* diffusion/kurtosis tensor, and *f* corresponds to the intra-axonal water fraction (Fieremans et al., 2011). As an illustrative supplementary analysis, we inverted the assumed relationship between diffusivities (intra-greater than extra-axonal), as follows:

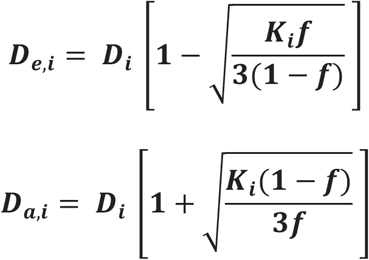

Because the model presumes aligned fibers and derives AWF from the maximum directional kurtosis (ignoring axonal dispersion and omitting intra-axonal diffusivity from the calculation, following Fieremans et al., 2011), AWF estimates are not affected by assignment of relative diffusivities. Conceptually, inverting the relative diffusivities essentially relabels the ‘faster diffusing’ compartment as ‘intra-axonal’. Of course, D_a,3_ > D_e,3_ would only be physically plausible in cases where, for example, axonal radii are large relative to the diffusion length, or where axonal dispersion is high and under sampled by the applied diffusion gradients; or perhaps where unanticipated signal exchange occurs across compartments. If D_a,3_ represents the direction roughly perpendicular to aligned axons (and if there is a direction for which D_a_ is approximately zero, i.e., diffusion is fully restricted), then D_a,3_ > D_e,3_ makes little sense. We are not arguing for the validity of the inverted model but rather hoping to explore the consequences of this specific change. Our illustrative analysis probably understates the complexity of the issue at hand, in part because we approximate AWF from maximal directional kurtosis, meaning that AWF estimates are not affected (Fieremans et al., 2011).

All values were mapped onto fiber tracts identified for each participant using the Automated Fiber Quantification software package (Yeatman et al., 2012b), freely available code written in MATLAB (https://github.com/yeatmanlab/AFQ), after initial generation of a whole-brain fiber estimates using the IFOD2 algorithm for probabilistic tractography in MRtrix version 3.0 (Tournier et al., 2019). Since the white matter tract integrity (WMTI) assumes that fibers are relatively well aligned (Fieremans et al., 2011), we followed recommendations from previous work and restricted our analysis to voxels with fractional anisotropy values greater than 0.3 (Chung et al., 2018a; Jensen et al., 2017a; Jensen et al., 2017b). Voxels with fractional anisotropy below 0.3 were removed and WMTI metrics were interpolated at each point on each fiber, and values were then summarized along the fiber-tract core by computing the median value across fiber nodes. Data with outlying values (greater than 4 standard deviations from the sample mean) in the white matter for any of the fitted metrics was excluded from further analysis. After excluding both outliers and individuals with excessive motion (>3mm, see above), the final data set included 111 sessions from 32 intervention participants and 75 sessions from 41 non-intervention control participants. A total of four control and five intervention participants were excluded at Session 1, four control and six intervention participants at Session 2, two control and three intervention participants at Session 3, and two control and four intervention participants at Session 4.

### Statistical Analysis

Statistical analysis was carried out using custom software written in MATLAB (version R2018a; https://github.com/yeatmanlab/Huber_2021_NeuroImage). To assess change over the course of intervention, we first averaged the middle 80% of each tract to create a single estimate of each property for each participant and tract. We selected the middle portion to eliminate the influence of crossing fibers near cortical terminations, and to avoid potential partial volume effects at the white matter / gray matter border. Mean tract values were then entered into a linear mixed effects model, with fixed effects of intervention time and a random effect of participant (random intercept). For quantifying intervention effects, we prefer to use ‘hours of intervention’, since this variable directly reflects the intervention ‘dose’. For analyses including the control participants, who did not participate in an intervention of any kind, we substitute ‘days since the baseline session’ for ‘hours of intervention’. Since sessions were held at regular intervals, the two variables were highly correlated (Pearson’s *r* = 0.94, *p* < 0.001).

## Results

### Diffusion MRI

Significant changes in mean diffusivity (MD) were apparent throughout the white matter in the intervention group (**Figure 3b, d;** note that raw coefficients are presented for descriptive purposes, while tracts showing significant change, qFDR < 0.05, are labeled accordingly). In a group of age-matched control participants who attended school as usual, we found no significant changes in white matter diffusivity over the same time frame, and growth estimates for the 16 pathways were distributed around zero (**Figure 3c**). A group (intervention vs. non-intervention) by time (days) interaction was significant (uncorrected p < 0.05) for the left arcuate fasciculus (*t*(182) = −1.97, *p* = 0.049).

**Figure 3.**
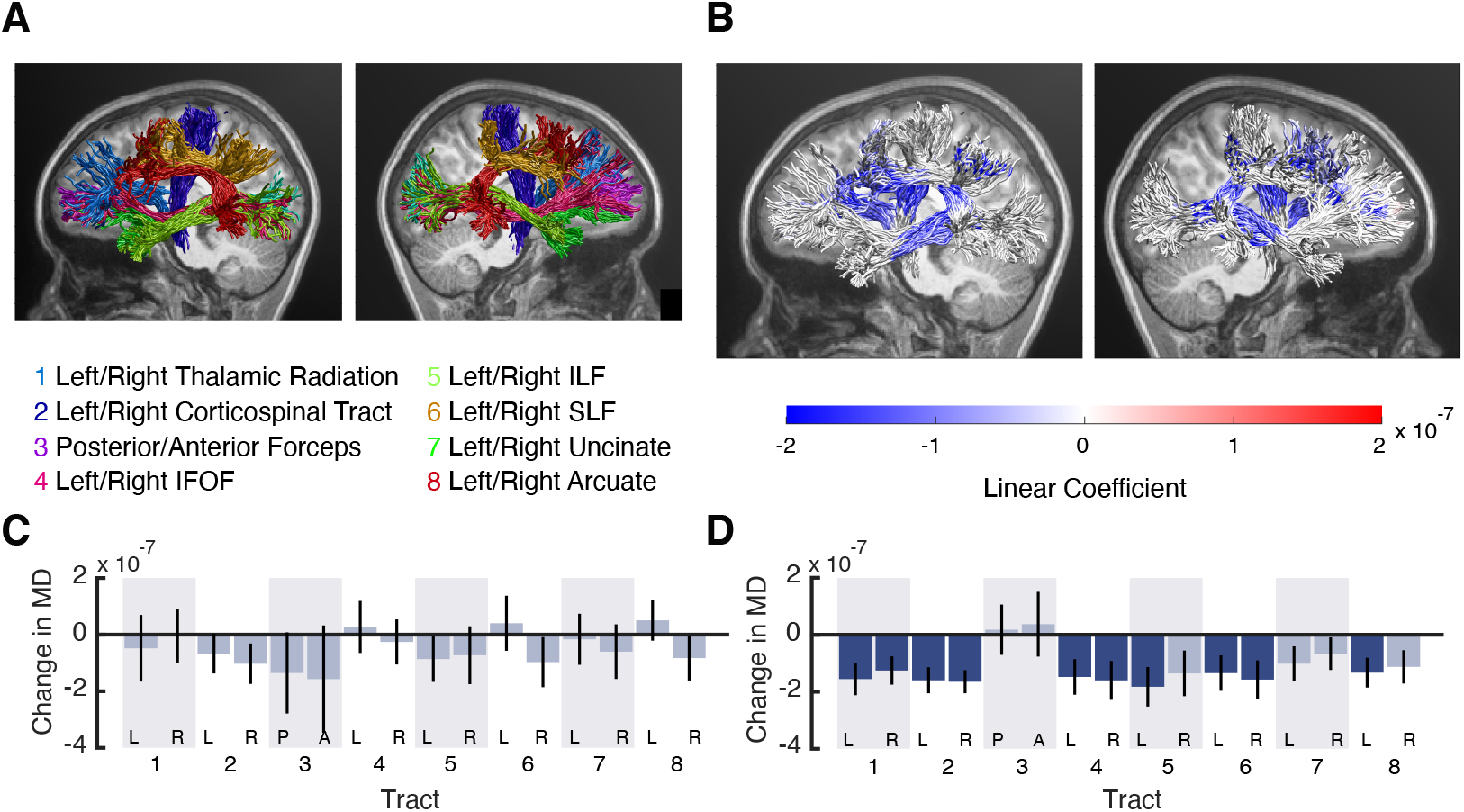
(A) White matter tracts are shown in two views (left hemisphere, right hemisphere) for an example participant. (B) The same tracts shown in (A), with color-coding based on the coefficient from a linear mixed effects model predicting mean diffusivity (MD) values as a function of intervention hours at each location along each tract. (C) Standardized coefficients from a mixed effects model (+/− 1 SE) predicting mean diffusivity from ‘days since the baseline session’, for the control group. No tracts showed significant effects, and estimates were distributed around zero. (D) Results for the intervention participants, using the same statistical model as in (C). Darker shaded bars reflect significant change (qFDR < 0.05).

Although mean diffusivity is a highly sensitive measure, it is not biologically specific (De Santis et al., 2014). We next examined the effects of intervention on parameters estimated from the WMTI model: axonal water fraction (AWF), intra-axonal diffusivity (Da), and extra-axonal mean diffusivity (MDe). As shown in **Figure 4**, intervention effects were limited to parameters associated with the extra-axonal space: Extra-axonal MD effects mirror the MD effects shown above. We saw no change in estimates of AWF over the intervention period. A supplementary analysis of axial, radial, and mean kurtosis gave identical results, with the exception of the right corticospinal tract, where a trend was present for mean (group-by-time interaction *t*(182) = −2.30, *p* = 0.023) and radial kurtosis (group-by-time interaction *t*(182) = −2.28, *p* = 0.024). No other tracts showed significant change in kurtosis estimates over the intervention period.

**Figure 4.**
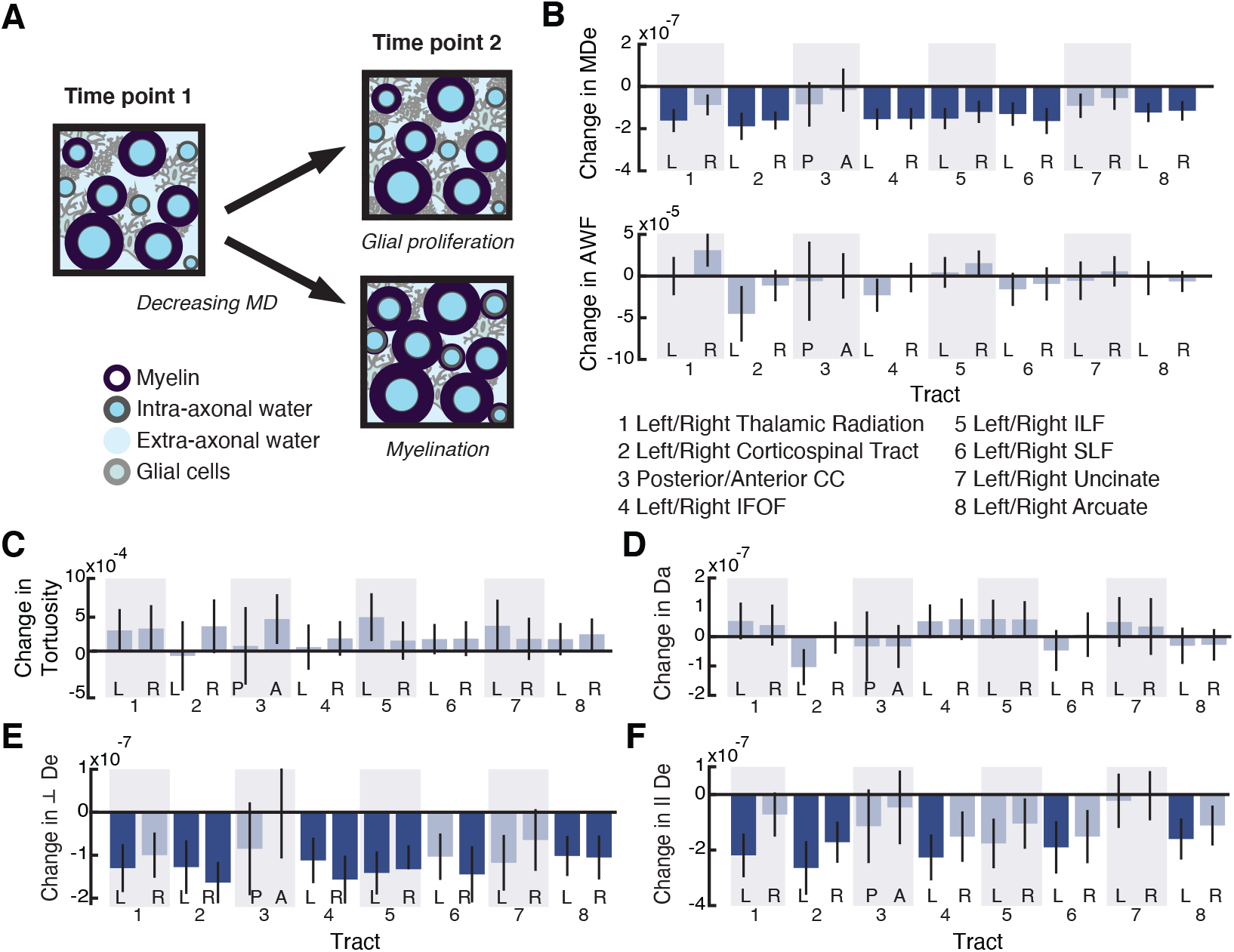
Modeling white matter plasticity. (A) Illustration of two scenarios in which mean diffusivity might decline in a voxel: proliferation of glial cells within the extra axonal space (top) or increasing axonal myelination (bottom). (B-F) Plots show coefficients from a linear mixed effects model predicting (B) extra-axonal mean diffusivity (MDe) and axon water fraction (AWF), (C) tortuosity 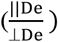, (D) intra-axonal diffusivity (Da), (E) perpendicular extra-axonal diffusivity (⊥De), and (F) parallel extra-axonal diffusivity (∥De) from intervention time in hours. Tracts showing significant change (*qFDR* < 0.05) are shaded with a darker blue.

Inverting the assumed relationship between diffusivities resulted in significant longitudinal effects attributed to intra-axonal diffusivity, as shown in **Figure 5,** and rendered the extra-axonal effects non-significant. We address the implications of this observation in the Discussion. Importantly, this change to the model does not affect our calculation of AWF (Fieremans et al., 2011), and so under either set of assumptions, we find no detectable difference in AWF over the course of the intervention.

**Figure 5.**
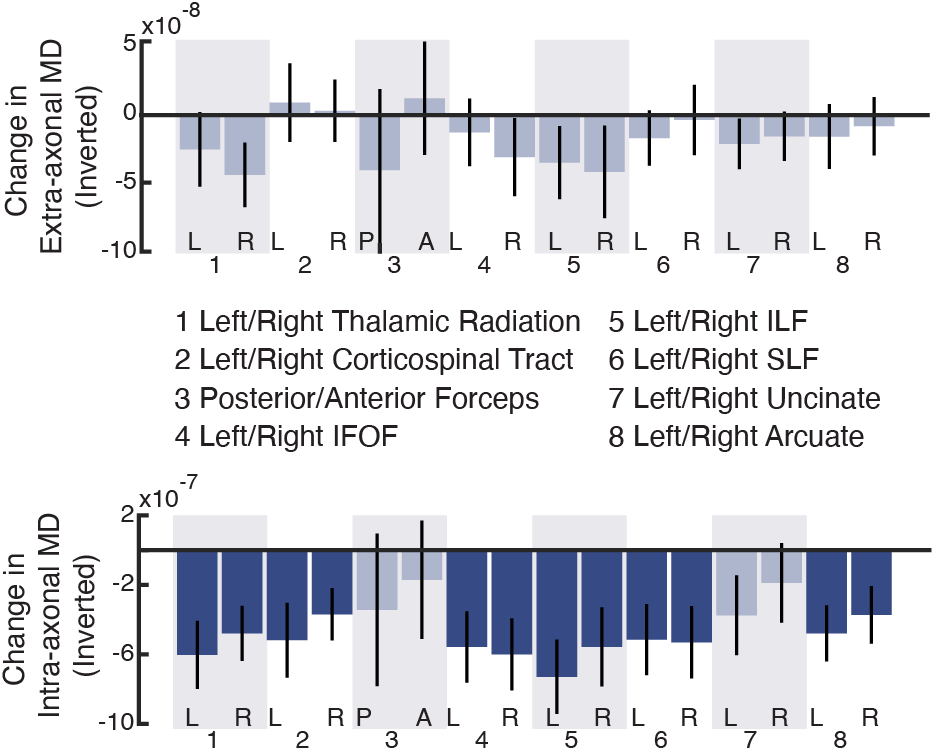
Inverting the assumed relationship between diffusivities rendered the extra-axonal effects non-significant (top) and resulted in significant longitudinal effects attributed to intra-axonal diffusivity (bottom). Plots show coefficients from a linear mixed effects model predicting each parameter from intervention time in hours. Tracts showing significant change (*qFDR* < 0.05) are shaded with a darker blue.

### Quantitative T1

The diffusion MRI effects are compatible with a process that influences diffusion within the extra-axonal space without changing the relative size of the axonal compartment. As discussed above, however, the WMTI parameters provide neither a direct nor unambiguos index of AWF. We therefore use qMRI data collected in the same participants during the intervention to further examine potential tissue changes. R1 measurements reflect both the total volume of tissue in a region, and the macro-molecular composition of that tissue, incuding lipid and iron content (Filo et al., 2019; Mezer et al., 2013; Stuber et al., 2014). A decrease in the volume of myelinated tissue (or, an increase in water) within a voxel would be associated with lower measured R1 values, while an increase in myelinated tissue would be associated with elevated R1 measurements. However, we found no significant change in R1 over the intervention period in either group, as shown in **Figure 6**. Baseline R1 values in severeal tracts did correlate with reading skill prior to the start of intervention (*p* < 0.05, uncorrected, see **Figure 6** for exact correlation values), as did MD and ‘extra-axonal’ MD values. The anatomical distribution of behavioral effects differed across modalities and parameters (R1 vs. WMTI derived diffusivities vs. AWF).

**Figure 6.**
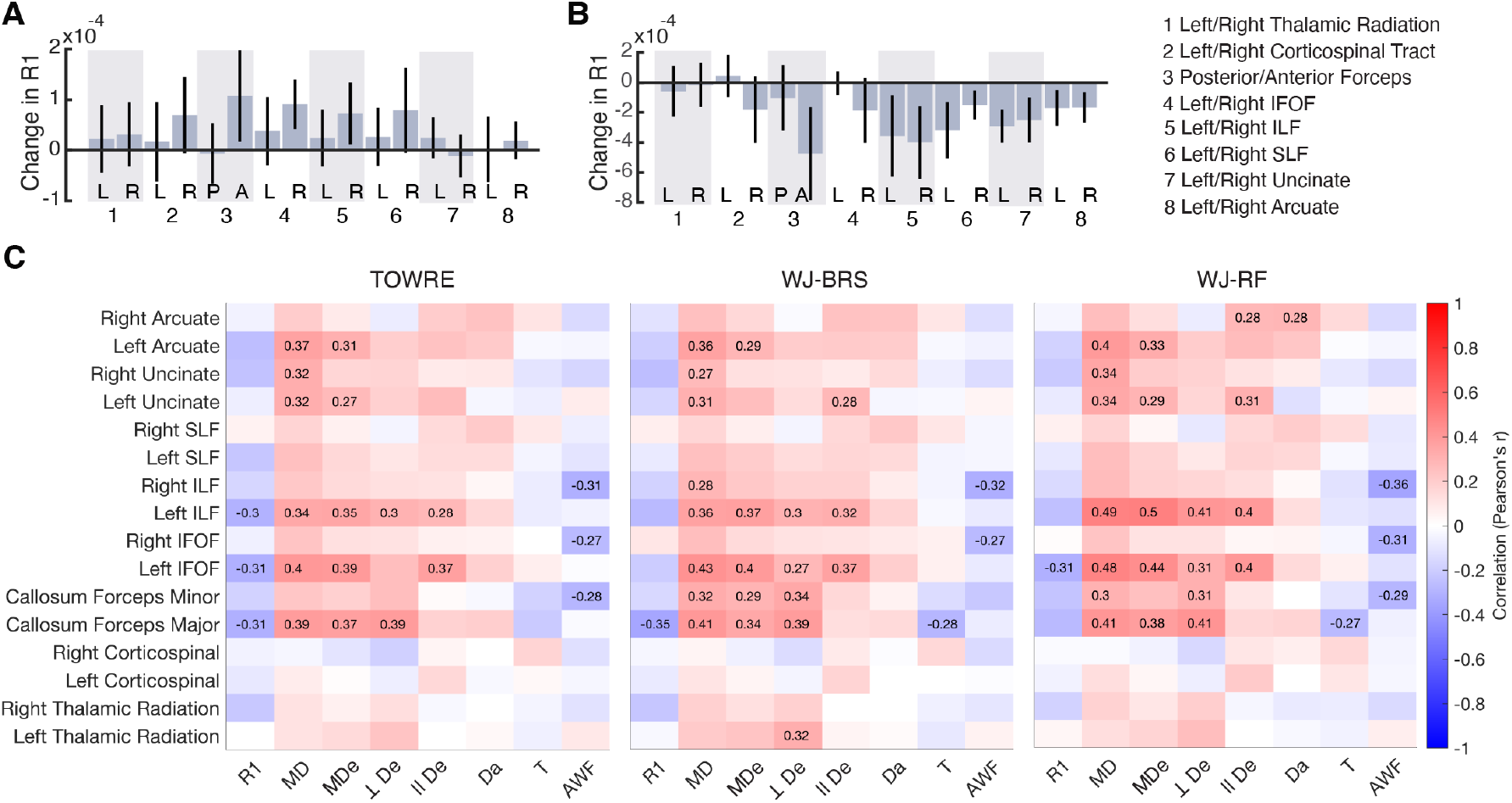
Quantitative R1 values show no significant change in the intervention group (A), or the non-intervention control group (B). Bar plots show coefficients from a linear mixed effects model predicting R1 from intervention time (days since baseline; random effect of participant (intercept)). (C) Correlations between baseline (pre-intervention) reading skill and baseline, tract averaged R1, MD, extra-axonal MD (MDe), extra-axonal perpendicular diffusivity (⊥De), extra-axonal axial diffusivity (∥De), intra-axonal diffusivity (Da), tortuosity (T), and axonal water fraction (AWF). Results are shown separately for each reading measure: The Test of Word Reading Efficiency index (TOWRE, far left), the Woodcock Johnson Tests of Achievement subtests Basic Reading Skill subtest (WJ-BRS, middle), and the Woodcock Johnson Reading Fluency subtest (WJ-RF, right). Color coding reflects individual Pearson’s correlation coefficients, and numerical values are given for all significant (*p* < 0.05, uncorrected) correlations.

## Discussion

Here we use diffusion MRI and a tissue model derived from diffusion kurtosis imaging (Fieremans et al., 2011), alongside quantitative R1 (1/T1 relaxation time) measurements, to examine changes in the white matter during a successful reading intervention. In both the intervention and non-intervention control groups, R1, MD, AWF, and ‘extra-axonal’ diffusivity correlated with pre-intervention reading skill and showed distinct anatomical distributions. While we observed systematic changes over the intervention in MD and ‘extra-axonal’ MD, we failed to see a change in estimated intra-axonal water fraction (AWF), or in quantitative R1 values.

While an absence of longitudinal change in either AWF or quantitative R1 points away from an underlying change in myelination, this null result should be interpreted cautiously. WMTI derived parameters have been evaluated in the context of both animal and human models of white matter pathology (Benitez et al., 2014; de Kouchkovsky et al., 2016; Falangola et al., 2014; Guglielmetti et al., 2016; Jelescu et al., 2015; Jelescu et al., 2016b; Kamiya et al., 2017; Kelm et al., 2016). AWF has been proposed as a marker for pathological axonal loss (Fieremans et al., 2013), but it has also been shown to vary with experimentally induced demyelination (Falangola et al., 2014; Jelescu et al., 2016b). Although diffusion MRI is not directly sensitive to myelin, it has been suggested that AWF might capture changes in myelination indirectly, if an associated change in the extra-axonal volume fraction were to alter the signal contributions from axonal versus non-axonal water (Jelescu et al., 2015; Jelescu et al., 2016b). However, it is not clear that the resulting signal change would be proportionate to the physical change in myelin. In contrast, R1 has been shown to vary systematically with myelin content, at least in the white matter (Stuber et al., 2014), and quantitative R1 estimates have been used to characterize age-related variation in white matter development (e.g., Yeatman et al., 2014). However, experience-driven changes in myelin might be still smaller or more variable across fibers than developmental or age-related effects. The complementary results for AWF and R1 thus provide converging information that argues against a widespread change in myelination, but do not unequivocally rule out the possibility that local myelin remodeling may be occurring.

We previously (Huber et al., 2018) reported changes in mean diffusivity alongside growth in reading skills during an intensive reading intervention. The current analysis suggests that these effects reflect properties of the extra-axonal space, rather than changes in myelination. It is plausible that a systematic and specific reduction in extra-axonal diffusivity could result from changes in the size or distribution of non-neuronal cells, such glia (Yi et al., 2019). This has been suggested elsewhere with respect to gray matter diffusivity (Blumenfeld-Katzir et al., 2011; Johansen-Berg et al., 2012; Sagi et al., 2012). In the white matter, developmental studies suggest that neuronal activity can affect the number of oligodendrocyte progenitor cells (OPCs) and potentially determine whether existing myelin sheaths are maintained (Hines et al., 2015). In adults, newborn OPCs are thought to participate in myelin remodeling and re-myelination, and animal models blocking OPC production show predicted impairments in motor skill learning (McKenzie et al., 2014; Xiao et al., 2016). Rapid OPC proliferation has been shown to precede later learning-related myelin remodeling (Gibson et al., 2014), with the early proliferative response appearing to exceed subsequent changes in myelination, perhaps reflecting an initial over-production of glial cells during an early stage of the learning process. Correlations between motor learning and diffusion indices associated with the untrained hemisphere have also been reported (Sampaio-Baptista et al., 2013), suggesting that diffusion measurements may capture spatially widespread variation relevant for learning, beyond the specific myelination of functionally relevant circuits.

It is important to consider the limitations of the diffusion MRI modeling framework used in the current study. The WMTI implementation used here assumes well-aligned fibers as its inputs, and we have tried to assure that this assumption is met in our analysis by sampling voxels with fractional anisotropy values that fall within a range for which the model has been applied (Chung et al., 2018a; Jensen et al., 2017b). It should be noted, however, that an FA cutoff of 0.3 likely includes the bulk of white matter, including regions with a variety of underlying geometries. As an illustrative example, we contrasted fits from the standard WMTI model (Fieremans et al., 2011) with fits from an implementation that assumed greater intra-versus extra-axonal diffusivity throughout the brain. As we note above, D_a, ⊥_ > D_e, ⊥_ would only be physically plausible in cases where, for example, axonal radii are large relative to the diffusion length, or where axonal dispersion is high and undersampled by the applied diffusion gradients (i.e., D_a, ⊥_ > 0 and/or does not correspond to diffusion perpendicular to aligned axons), or perhaps where unanticipated signal exchange occurs across compartments. Whether overall D_a_ is in fact greater than D_e, ⊥_ remains a topic of continued study (see Jelescu et al., 2020 for recent review; see also Jespersen et al., 2018; Kunz et al., 2018). Here, we observed that intervention effects were localized to the intra-rather than extra-axonal space when the relative diffusivities were inverted. Alteration in intra-axonal diffusivity has previously been tied to axonal injury (Chung et al., 2018b), although it is less clear what non-pathological mechanism would alter the intrinsic diffusivity of the intra-axonal space over the timescales and in the participant sample considered here. Nonetheless, it is worth noting that the biological interpretation of these effects depends critically on the derivation of compartment diffusivities within the model. Our illustrative analysis probably understates the complexity of the issue at hand, in part because we approximate AWF from maximal directional kurtosis and therefore our estimates of AWF are less biologically exact, but stable (Fieremans et al., 2011). While there has been considerable recent effort aimed at establishing more flexible models that are biologically valid and computationally robust, substantial work remains (see Jelescu et al., 2020 for recent review).

Consistent with a number of previous dMRI studies in children (Deutsch et al., 2005; Lebel et al., 2013; Odegard et al., 2009; Travis et al., 2017; Vanderauwera et al., 2018; Yeatman et al., 2012a; Yeatman et al., 2011), we found a correlation between diffusion indices and reading skill within tracts considered core to the reading and language circuitry (Ben-Shachar et al., 2007; Vandermosten et al., 2012; Wandell and Yeatman, 2013), namely, the arcuate fasciculus, uncinate, ILF, IFOF, and the posterior and anterior crossing of the corpus collosum. Correlations with quantitative R1 estimates highlighted partly overlapping anatomical regions, specifically, the left ILF and IFOF and the posterior callosal fibers. Lower estimated AWF in the right ILF, IFOF, and the anterior callosal fibers also predicted higher reading skill. The analysis of behavioral correlations was exploratory in nature, and not a primary aim of the current paper; however, it underscores the notion that distinct mechanisms may underpin correlations between specific white matter connections and individual differences in reading over development, a topic which merits further study.

The relationship between higher-level cognitive function and structural features of myelinated and unmyelinated neurites, glial cells, and vasculature is surely complex, and the process of maintaining and optimizing this tissue architecture likely involves several distinct biological phenomena that operate over different time scales. Maturational differences in diffusion within the white matter reflect changes in myelination that occur over the timescale of years (Chang et al., 2015; Jelescu et al., 2015). The current data suggest that short-term changes in diffusion properties might capture an initial stage of the learning process, although the connection between rapid non-neuronal changes and long-term remodeling of myelin is not fully resolved. Studies employing longitudinal measurements, coupled with increasingly sophisticated imaging and modeling techniques, hold promise for revealing the interplay among distinct biological processes that support learning and cognition. Understanding the relationship between learning and plasticity at temporal scales ranging from hours (Hofstetter et al.; Hofstetter et al.; Sagi et al., 2012), to days (Huber et al., 2019), to years (Wang et al., 2017; Yeatman et al., 2012a), will require research aimed at forging a tighter link between education and neuroscience.

## Acknowledgements

This work was funded by NSF/BSF BCS #1551330/ #2015608, Eunice Kennedy Shriver National Institute of Child Health and Human Development Grant R01 HD09586101 and Jacobs Foundation Research Fellowship to JDY.

## References

Alexander, D.C., Dyrby, T.B., Nilsson, M., Zhang, H., 2019. Imaging brain microstructure with diffusion MRI: practicality and applications. NMR Biomed 32, e3841.

Andersson, J.L., Skare, S., Ashburner, J., 2003. How to correct susceptibility distortions in spin-echo echo-planar images: application to diffusion tensor imaging. Neuroimage 20, 870–888.

Andersson, J.L., Sotiropoulos, S.N., 2016. An integrated approach to correction for off-resonance effects and subject movement in diffusion MR imaging. Neuroimage 125, 1063–1078.

Ashburner, J., Friston, K., 1997. Multimodal image coregistration and partitioning--a unified framework. Neuroimage 6, 209–217.

Barral, J.K., Gudmundson, E., Stikov, N., Etezadi-Amoli, M., Stoica, P., Nishimura, D.G., 2010. A robust methodology for in vivo T1 mapping. Magn Reson Med 64, 1057–1067.

Basser, P.J., Mattiello, J., LeBihan, D., 1994. Estimation of the effective self-diffusion tensor from the NMR spin echo. J Magn Reson B 103, 247–254.

Bechler, M.E., Swire, M., Ffrench-Constant, C., 2018. Intrinsic and adaptive myelination-A sequential mechanism for smart wiring in the brain. Dev Neurobiol 78, 68–79.

Bell, N., 2007. Seeing stars. Gander, San Luis Obispo, CA.

Ben-Shachar, M., Dougherty, R.F., Wandell, B.A., 2007. White matter pathways in reading. Curr Opin Neurobiol 17, 258–270.

Benitez, A., Fieremans, E., Jensen, J.H., Falangola, M.F., Tabesh, A., Ferris, S.H., Helpern, J.A., 2014. White matter tract integrity metrics reflect the vulnerability of late-myelinating tracts in Alzheimer’s disease. Neuroimage Clin 4, 64–71.

Blumenfeld-Katzir, T., Pasternak, O., Dagan, M., Assaf, Y., 2011. Diffusion MRI of structural brain plasticity induced by a learning and memory task. PLoS One 6, e20678.

Cercignani, M., Bouyagoub, S., 2018. Brain microstructure by multi-modal MRI: Is the whole greater than the sum of its parts? Neuroimage 182, 117–127.

Chang, Y.S., Owen, J.P., Pojman, N.J., Thieu, T., Bukshpun, P., Wakahiro, M.L., Berman, J.I., Roberts, T.P., Nagarajan, S.S., Sherr, E.H., Mukherjee, P., 2015. White Matter Changes of Neurite Density and Fiber Orientation Dispersion during Human Brain Maturation. PLoS One 10, e0123656.

Chung, S., Fieremans, E., Kucukboyaci, N.E., Wang, X., Morton, C.J., Novikov, D.S., Rath, J.F., Lui, Y.W., 2018a. Working Memory And Brain Tissue Microstructure: White Matter Tract Integrity Based On Multi-Shell Diffusion MRI. Sci Rep 8, 3175.

Chung, S., Fieremans, E., Wang, X., Kucukboyaci, N.E., Morton, C.J., Babb, J., Amorapanth, P., Foo, F.A., Novikov, D.S., Flanagan, S.R., Rath, J.F., Lui, Y.W., 2018b. White Matter Tract Integrity: An Indicator of Axonal Pathology after Mild Traumatic Brain Injury. J Neurotrauma 35, 1015–1020.

de Kouchkovsky, I., Fieremans, E., Fleysher, L., Herbert, J., Grossman, R.I., Inglese, M., 2016. Quantification of normal-appearing white matter tract integrity in multiple sclerosis: a diffusion kurtosis imaging study. J Neurol 263, 1146–1155.

De Santis, S., Drakesmith, M., Bells, S., Assaf, Y., Jones, D.K., 2014. Why diffusion tensor MRI does well only some of the time: variance and covariance of white matter tissue microstructure attributes in the living human brain. Neuroimage 89, 35–44.

Deutsch, G.K., Dougherty, R.F., Bammer, R., Siok, W.T., Gabrieli, J.D., Wandell, B., 2005. Children’s reading performance is correlated with white matter structure measured by diffusion tensor imaging. Cortex 41, 354–363.

Ekerdt, C.E.M., Kuhn, C., Anwander, A., Brauer, J., Friederici, A.D., 2020. Word learning reveals white matter plasticity in preschool children. Brain Struct Funct 225, 607–619.

Falangola, M.F., Guilfoyle, D.N., Tabesh, A., Hui, E.S., Nie, X., Jensen, J.H., Gerum, S.V., Hu, C., LaFrancois, J., Collins, H.R., Helpern, J.A., 2014. Histological correlation of diffusional kurtosis and white matter modeling metrics in cuprizone-induced corpus callosum demyelination. NMR Biomed 27, 948–957.

Fields, R.D., 2015. A new mechanism of nervous system plasticity: activity-dependent myelination. Nat Rev Neurosci 16, 756–767.

Fieremans, E., Benitez, A., Jensen, J.H., Falangola, M.F., Tabesh, A., Deardorff, R.L., Spampinato, M.V., Babb, J.S., Novikov, D.S., Ferris, S.H., Helpern, J.A., 2013. Novel white matter tract integrity metrics sensitive to Alzheimer disease progression. AJNR Am J Neuroradiol 34, 2105–2112.

Fieremans, E., Jensen, J.H., Helpern, J.A., 2011. White matter characterization with diffusional kurtosis imaging. Neuroimage 58, 177–188.

Filo, S., Shtangel, O., Salamon, N., Kol, A., Weisinger, B., Shifman, S., Mezer, A.A., 2019. Disentangling molecular alterations from water-content changes in the aging human brain using quantitative MRI. Nat Commun 10, 3403.

Garyfallidis, E., Brett, M., Amirbekian, B., Rokem, A., van der Walt, S., Descoteaux, M., Nimmo-Smith, I., Dipy, C., 2014. Dipy, a library for the analysis of diffusion MRI data. Front Neuroinform 8, 8.

Geeraert, B.L., Lebel, R.M., Lebel, C., 2019. A multiparametric analysis of white matter maturation during late childhood and adolescence. Hum Brain Mapp 40, 4345–4356.

Genc, S., Malpas, C.B., Holland, S.K., Beare, R., Silk, T.J., 2017. Neurite density index is sensitive to age related differences in the developing brain. Neuroimage 148, 373–380.

Gibson, E.M., Purger, D., Mount, C.W., Goldstein, A.K., Lin, G.L., Wood, L.S., Inema, I., Miller, S.E., Bieri, G., Zuchero, J.B., Barres, B.A., Woo, P.J., Vogel, H., Monje, M., 2014. Neuronal activity promotes oligodendrogenesis and adaptive myelination in the mammalian brain. Science 344, 1252304.

Guglielmetti, C., Veraart, J., Roelant, E., Mai, Z., Daans, J., Van Audekerke, J., Naeyaert, M., Vanhoutte, G., Delgado, Y.P.R., Praet, J., Fieremans, E., Ponsaerts, P., Sijbers, J., Van der Linden, A., Verhoye, M., 2016. Diffusion kurtosis imaging probes cortical alterations and white matter pathology following cuprizone induced demyelination and spontaneous remyelination. Neuroimage 125, 363–377.

Henriques Rafael Neto, C.M.M., Marrale Maurizio, Huber Elizabeth, Kruper John, Koudoro Serge, Yeatman Jason D., Garyfallidis Eleftherios, Rokem Ariel, 2021. Diffusional Kurtosis Imaging in the Diffusion Imaging in Python Project. Frontiers in Human Neuroscience 15, 390.

Hines, J.H., Ravanelli, A.M., Schwindt, R., Scott, E.K., Appel, B., 2015. Neuronal activity biases axon selection for myelination in vivo. Nat Neurosci 18, 683–689.

Hofstetter, S., Friedmann, N., Assaf, Y., 2017. Rapid language-related plasticity: microstructural changes in the cortex after a short session of new word learning. Brain Struct Funct 222, 1231–1241.

Hofstetter, S., Tavor, I., Tzur Moryosef, S., Assaf, Y., 2013. Short-term learning induces white matter plasticity in the fornix. J Neurosci 33, 12844–12850.

Huber, E., Donnelly, P.M., Rokem, A., Yeatman, J.D., 2018. Rapid and widespread white matter plasticity during an intensive reading intervention. Nat Commun 9, 2260.

Huber, E., Henriques, R.N., Owen, J.P., Rokem, A., Yeatman, J.D., 2019. Applying microstructural models to understand the role of white matter in cognitive development. Dev Cogn Neurosci 36, 100624.

Jelescu, I.O., Budde, M.D., 2017. Design and validation of diffusion MRI models of white matter. Front Phys 28.

Jelescu, I.O., Palombo, M., Bagnato, F., Schilling, K.G., 2020. Challenges for biophysical modeling of microstructure. J Neurosci Methods 344, 108861.

Jelescu, I.O., Veraart, J., Adisetiyo, V., Milla, S.S., Novikov, D.S., Fieremans, E., 2015. One diffusion acquisition and different white matter models: how does microstructure change in human early development based on WMTI and NODDI? Neuroimage 107, 242–256.

Jelescu, I.O., Veraart, J., Fieremans, E., Novikov, D.S., 2016a. Degeneracy in model parameter estimation for multi-compartmental diffusion in neuronal tissue. NMR Biomed 29, 33–47.

Jelescu, I.O., Zurek, M., Winters, K.V., Veraart, J., Rajaratnam, A., Kim, N.S., Babb, J.S., Shepherd, T.M., Novikov, D.S., Kim, S.G., Fieremans, E., 2016b. In vivo quantification of demyelination and recovery using compartment-specific diffusion MRI metrics validated by electron microscopy. Neuroimage 132, 104–114.

Jensen, A., Stickley, T., Torrissen, W., Stigmar, K., 2017a. Arts on prescription in Scandinavia: a review of current practice and future possibilities. Perspect Public Health 137, 268–274.

Jensen, J.H., Helpern, J.A., 2010. MRI quantification of non-Gaussian water diffusion by kurtosis analysis. NMR Biomed 23, 698–710.

Jensen, J.H., Helpern, J.A., Ramani, A., Lu, H., Kaczynski, K., 2005. Diffusional kurtosis imaging: the quantification of non-gaussian water diffusion by means of magnetic resonance imaging. Magn Reson Med 53, 1432–1440.

Jensen, J.H., McKinnon, E.T., Glenn, G.R., Helpern, J.A., 2017b. Evaluating kurtosis-based diffusion MRI tissue models for white matter with fiber ball imaging. NMR Biomed 30.

Jensen, J.H., Russell Glenn, G., Helpern, J.A., 2016. Fiber ball imaging. Neuroimage 124, 824–833.

Jespersen, S.N., Bjarkam, C.R., Nyengaard, J.R., Chakravarty, M.M., Hansen, B., Vosegaard, T., Ostergaard, L., Yablonskiy, D., Nielsen, N.C., Vestergaard-Poulsen, P., 2010. Neurite density from magnetic resonance diffusion measurements at ultrahigh field: comparison with light microscopy and electron microscopy. Neuroimage 49, 205–216.

Jespersen, S.N., Olesen, J.L., Hansen, B., Shemesh, N., 2018. Diffusion time dependence of microstructural parameters in fixed spinal cord. Neuroimage 182, 329–342.

Johansen-Berg, H., Baptista, C.S., Thomas, A.G., 2012. Human structural plasticity at record speed. Neuron 73, 1058–1060.

Jolles, D., Wassermann, D., Chokhani, R., Richardson, J., Tenison, C., Bammer, R., Fuchs, L., Supekar, K., Menon, V., 2016. Plasticity of left perisylvian white-matter tracts is associated with individual differences in math learning. Brain Struct Funct 221, 1337–1351.

Kaden, E., Kelm, N.D., Carson, R.P., Does, M.D., Alexander, D.C., 2016. Multi-compartment microscopic diffusion imaging. Neuroimage 139, 346–359.

Kamiya, K., Hori, M., Irie, R., Miyajima, M., Nakajima, M., Kamagata, K., Tsuruta, K., Saito, A., Nakazawa, M., Suzuki, Y., Mori, H., Kunimatsu, A., Arai, H., Aoki, S., Abe, O., 2017. Diffusion imaging of reversible and irreversible microstructural changes within the corticospinal tract in idiopathic normal pressure hydrocephalus. Neuroimage Clin 14, 663–671.

Kelm, N.D., West, K.L., Carson, R.P., Gochberg, D.F., Ess, K.C., Does, M.D., 2016. Evaluation of diffusion kurtosis imaging in ex vivo hypomyelinated mouse brains. Neuroimage 124, 612–626.

Kraft, I., Schreiber, J., Cafiero, R., Metere, R., Schaadt, G., Brauer, J., Neef, N.E., Muller, B., Kirsten, H., Wilcke, A., Boltze, J., Friederici, A.D., Skeide, M.A., 2016. Predicting early signs of dyslexia at a preliterate age by combining behavioral assessment with structural MRI. Neuroimage 143, 378–386.

Kunz, N., da Silva, A.R., Jelescu, I.O., 2018. Intra- and extra-axonal axial diffusivities in the white matter: Which one is faster? Neuroimage 181, 314–322.

Lebel, C., Beaulieu, C., 2011. Longitudinal development of human brain wiring continues from childhood into adulthood. J Neurosci 31, 10937–10947.

Lebel, C., Shaywitz, B., Holahan, J., Shaywitz, S., Marchione, K., Beaulieu, C., 2013. Diffusion tensor imaging correlates of reading ability in dysfluent and non-impaired readers. Brain Lang 125, 215–222.

Mamiya, P.C., Richards, T.L., Coe, B.P., Eichler, E.E., Kuhl, P.K., 2016. Brain white matter structure and COMT gene are linked to second-language learning in adults. Proc Natl Acad Sci U S A 113, 7249–7254.

McKenzie, I.A., Ohayon, D., Li, H., de Faria, J.P., Emery, B., Tohyama, K., Richardson, W.D., 2014. Motor skill learning requires active central myelination. Science 346, 318–322.

Metzler-Baddeley, C., Foley, S., de Santis, S., Charron, C., Hampshire, A., Caeyenberghs, K., Jones, D.K., 2017. Dynamics of White Matter Plasticity Underlying Working Memory Training: Multimodal Evidence from Diffusion MRI and Relaxometry. J Cogn Neurosci 29, 1509–1520.

Mezer, A., Rokem, A., Berman, S., Hastie, T., Wandell, B.A., 2016. Evaluating quantitative proton-density-mapping methods. Hum Brain Mapp 37, 3623–3635.

Mezer, A., Yeatman, J.D., Stikov, N., Kay, K.N., Cho, N.J., Dougherty, R.F., Perry, M.L., Parvizi, J., Hua le, H., Butts-Pauly, K., Wandell, B.A., 2013. Quantifying the local tissue volume and composition in individual brains with magnetic resonance imaging. Nat Med 19, 1667–1672.

Moura, L.M., Kempton, M., Barker, G., Salum, G., Gadelha, A., Pan, P.M., Hoexter, M., Del Aquilla, M.A., Picon, F.A., Anes, M., Otaduy, M.C., Amaro, E., Jr., Rohde, L.A., McGuire, P., Bressan, R.A., Sato, J.R., Jackowski, A.P., 2016. Age-effects in white matter using associated diffusion tensor imaging and magnetization transfer ratio during late childhood and early adolescence. Magn Reson Imaging 34, 529–534.

Novikov, D.S., Veraart, J., Jelescu, I.O., Fieremans, E., 2018. Rotationally-invariant mapping of scalar and orientational metrics of neuronal microstructure with diffusion MRI. Neuroimage 174, 518–538.

Odegard, T.N., Farris, E.A., Ring, J., McColl, R., Black, J., 2009. Brain connectivity in non-reading impaired children and children diagnosed with developmental dyslexia. Neuropsychologia 47, 1972–1977.

Sagi, Y., Tavor, I., Hofstetter, S., Tzur-Moryosef, S., Blumenfeld-Katzir, T., Assaf, Y., 2012. Learning in the fast lane: new insights into neuroplasticity. Neuron 73, 1195–1203.

Sampaio-Baptista, C., Johansen-Berg, H., 2017. White Matter Plasticity in the Adult Brain. Neuron 96, 1239–1251.

Sampaio-Baptista, C., Khrapitchev, A.A., Foxley, S., Schlagheck, T., Scholz, J., Jbabdi, S., DeLuca, G.C., Miller, K.L., Taylor, A., Thomas, N., Kleim, J., Sibson, N.R., Bannerman, D., Johansen-Berg, H., 2013. Motor skill learning induces changes in white matter microstructure and myelination. J Neurosci 33, 19499–19503.

Stuber, C., Morawski, M., Schafer, A., Labadie, C., Wahnert, M., Leuze, C., Streicher, M., Barapatre, N., Reimann, K., Geyer, S., Spemann, D., Turner, R., 2014. Myelin and iron concentration in the human brain: a quantitative study of MRI contrast. Neuroimage 93 Pt 1, 95–106.

Takemura, H., Ogawa, S., Mezer, A.A., Horiguchi, H., Miyazaki, A., Matsumoto, K., Shikishima, K., Nakano, T., Masuda, Y., 2019. Diffusivity and quantitative T1 profile of human visual white matter tracts after retinal ganglion cell damage. Neuroimage Clin 23, 101826.

Tamnes, C.K., Roalf, D.R., Goddings, A.L., Lebel, C., 2018. Diffusion MRI of white matter microstructure development in childhood and adolescence: Methods, challenges and progress. Dev Cogn Neurosci 33, 161–175.

Tournier, J.D., 2019. Diffusion MRI in the brain - Theory and concepts. Prog Nucl Magn Reson Spectrosc 112-113, 1–16.

Tournier, J.D., Smith, R., Raffelt, D., Tabbara, R., Dhollander, T., Pietsch, M., Christiaens, D., Jeurissen, B., Yeh, C.H., Connelly, A., 2019. MRtrix3: A fast, flexible and open software framework for medical image processing and visualisation. Neuroimage 202, 116137.

Travis, K.E., Adams, J.N., Kovachy, V.N., Ben-Shachar, M., Feldman, H.M., 2017. White matter properties differ in 6-year old Readers and Pre-readers. Brain Struct Funct 222, 1685–1703.

Travis, K.E., Castro, M.R.H., Berman, S., Dodson, C.K., Mezer, A.A., Ben-Shachar, M., Feldman, H.M., 2019. More than myelin: Probing white matter differences in prematurity with quantitative T1 and diffusion MRI. Neuroimage Clin 22, 101756.

Tustison, N.J., Avants, B.B., Gee, J.C., 2009. Directly manipulated free-form deformation image registration. IEEE Trans Image Process 18, 624–635.

Vanderauwera, J., De Vos, A., Forkel, S.J., Catani, M., Wouters, J., Vandermosten, M., Ghesquiere, P., 2018. Neural organization of ventral white matter tracts parallels the initial steps of reading development: A DTI tractography study. Brain Lang 183, 32–40.

Vandermosten, M., Boets, B., Wouters, J., Ghesquiere, P., 2012. A qualitative and quantitative review of diffusion tensor imaging studies in reading and dyslexia. Neurosci Biobehav Rev 36, 1532–1552.

Wandell, B.A., Yeatman, J.D., 2013. Biological development of reading circuits. Curr Opin Neurobiol 23, 261–268.

Wang, Y., Mauer, M.V., Raney, T., Peysakhovich, B., Becker, B.L.C., Sliva, D.D., Gaab, N., 2017. Development of Tract-Specific White Matter Pathways During Early Reading Development in At-Risk Children and Typical Controls. Cereb Cortex 27, 2469–2485.

Xiao, L., Ohayon, D., McKenzie, I.A., Sinclair-Wilson, A., Wright, J.L., Fudge, A.D., Emery, B., Li, H., Richardson, W.D., 2016. Rapid production of new oligodendrocytes is required in the earliest stages of motor-skill learning. Nat Neurosci 19, 1210–1217.

Yeatman, J.D., Dougherty, R.F., Ben-Shachar, M., Wandell, B.A., 2012a. Development of white matter and reading skills. Proc Natl Acad Sci U S A 109, E3045–3053.

Yeatman, J.D., Dougherty, R.F., Myall, N.J., Wandell, B.A., Feldman, H.M., 2012b. Tract profiles of white matter properties: automating fiber-tract quantification. PLoS One 7, e49790.

Yeatman, J.D., Dougherty, R.F., Rykhlevskaia, E., Sherbondy, A.J., Deutsch, G.K., Wandell, B.A., Ben-Shachar, M., 2011. Anatomical properties of the arcuate fasciculus predict phonological and reading skills in children. J Cogn Neurosci 23, 3304–3317.

Yeatman, J.D., Wandell, B.A., Mezer, A.A., 2014. Lifespan maturation and degeneration of human brain white matter. Nat Commun 5, 4932.

Yendiki, A., Koldewyn, K., Kakunoori, S., Kanwisher, N., Fischl, B., 2014. Spurious group differences due to head motion in a diffusion MRI study. Neuroimage 88, 79–90.

Yi, S.Y., Barnett, B.R., Torres-Velazquez, M., Zhang, Y., Hurley, S.A., Rowley, P.A., Hernando, D., Yu, J.J., 2019. Detecting Microglial Density With Quantitative Multi-Compartment Diffusion MRI. Front Neurosci 13, 81.

Zhang, H., Schneider, T., Wheeler-Kingshott, C.A., Alexander, D.C., 2012. NODDI: practical in vivo neurite orientation dispersion and density imaging of the human brain. Neuroimage 61, 1000–1016.

